# Conformational Diversity and Interaction Signatures of NADH across protein families

**DOI:** 10.64898/2026.02.15.704522

**Authors:** Dubey Saniya, Chinmay Majee, Sumohana S. Channappayya, Eerappa Rajakumara

## Abstract

Nicotinamide adenine dinucleotide (NADH) is a ubiquitous redox cofactor that participates in a wide range of enzymatic and regulatory processes. These include metabolism, signalling, and diseases such as cancer and neurodegeneration. Despite the abundance of NADH–protein complex structures, general principles governing how proteins shape NADH conformation and interaction modes remain unclear, which limits our ability to rationally interpret cofactor specificity, catalytic efficiency, and off-target effects in inhibitors. Here, we present a comprehensive structural analysis of NADH recognition across protein families using 345 NADH-bound crystal structures from the Protein Data Bank. We adopted a descriptor-driven strategy that quantitatively captures the internal geometry of NADH using angles, dihedrals, and interatomic distances, enabling direct comparison of cofactor shapes independent of protein fold. These studies reveal that 65% of structures preferred conformers with a conserved adenine– nicotinamide separation while allowing limited flexibility in the pyrophosphate. The interaction profiles demonstrate that NADH recognition is dominated by hydrogen bonding and electrostatic interactions involving nearly all heteroatoms, while most carbon positions remain non-interacting. Residue and moiety level analyses further show that the nicotinamide region acts as the primary interaction hotspot across enzyme classes, while only a handful of structures exhibit adenine-centric recognition. Together, this study establishes a unified biophysical framework that links NADH shape, interaction signatures, and protein context, providing rational insights for cofactor engineering and the design of NADH-targeted inhibitors.

## Introduction

Nicotinamide Adenine Dinucleotide (NAD) is one of the universal cofactors in biology and exists in two interconvertible redox states, the oxidized form (NAD□) and the reduced form (NADH) [1]. While NAD+ primarily serves as an electron acceptor in oxidative pathways, NADH acts as an electron donor in reductive processes [2]. The balance of NAD+ and NADH is called the NAD+/NADH ratio [3]. NAD is one of the crucial metabolites in the cell. Its levels are essential for the normal physiology of the cell and are strictly regulated to prevent pathological conditions. NAD functions as a substrate of regulatory proteins involved in cell survival, DNA damage repair, calcium signaling, transcription regulation etc [4]. It also serves as a mediator of protein-protein interactions in various signaling pathways [5]. The major function of NADH and its oxidized counterpart (NAD^+^) is that of a redox couple that regulates a plethora of biological processes such as calcium homeostasis, synaptic plasticity, anti-apoptosis, and gene expression [6]. NAD^+^/NADH homeostasis is efficiently regulated, and any imbalance in levels of any of these results in mitochondrial dysfunction, which plays a crucial role in the cascade of various neurodegenerative disorders such as Alzheimer’s and Parkinson’s disease [6]. NAD-dependent enzymes include sirtuins (SIRTs), poly- and mono-(ADP-ribose) polymerases (PARPs and MARPs), cyclic ADP ribose hydrolases, NADH-dependent cytochrome p450 reductases, etc [7]. NADH-dependent oxidoreductases are a highly diverse group of enzymes that act on a vast range of substrates to catalyze electron transfer reactions. These are also important in a broad range of diverse regulatory and functional roles in human biology [8]. While NAD is popularly known as the intracellular redox factor, extracellular NAD can function as a signalling ligand by interacting with cell surface receptors or receptor-associated systems [9]. One of the examples of these is the purinergic receptor P2X7 in mouse immune cells, which is activated by NAD indirectly by ADP-ribosylation mediated by ectoenzymes such as ART, which leads to calcium influx, pore formation and cell death [10–12]. In some cases, NAD also acts on P2Y purinergic receptors, such as P2Y11, modulating intracellular cAMP and calcium signalling in immune and epithelial cells [13–15].

Numerous studies have explored the potential of NADH in treating and preventing various diseases. One of the most studied is Parkinson’s disease. The findings from these studies suggest that NADH shows therapeutic potential through multiple mechanisms, such as modulating neurotransmitter systems, enhancing ATP production, protecting against dopaminergic neuron degeneration, and reducing neuroinflammation [16]. One study puts forth a theory that NADH can be used as a medicine for cancer and can also combat the wastage and weakness of cancer patients [17]. NAD analogs, like FDA-approved drugs Olaparib and Rucaparib, work as PARP inhibitors by blocking Poly (ADP-Ribose) Polymerase (PARP) enzymes, which use NAD+ to repair DNA damage[18].

Compared to proteins, there are fewer biological ligands that exhibit conformational variability. It is clear that just a few rotatable bonds can lead to a large number of conformational possibilities for a ligand. The conformational freedom of a ligand is either restricted to a small locus, or in some cases there is retention of mobility after it binds to a protein [19–21]. The design of inhibitors targeting binding pockets for NAD is sometimes criticized on the basis that such inhibitors are likely to have off-target effects [22].

In this article, we have studied and analyzed 345 PDB entries containing NADH as the co-crystallized complex with enzymes. The structural aspects of the conserved, unique, and differentiating NADH interactions among various proteins have been studied to gain insights into and knowledge of NADH structural plasticity and functional promiscuity, which may pave the way for designing and developing effective and efficient lead compounds against various diseases.

## Materials and Methods

### Data Collection

To study the interactions of NADH across diverse proteins, a dataset of all PDB entries containing NADH in complex with proteins present in the RCSB-PDB repository was created using the advanced search option, with the filter key (feature) chemical ID set to ‘NADH’ and its value as chemical ID [23]. This chemical ID was the PubChem ID of NADH, which is 439153 [24]. This resulted in 345 structures. All the structures were downloaded in 8 different batches to make a library of the structures.

### Methodology For Descriptor Development

To get better insights, all the NADH structures extracted from 345 complexes were superimposed using the “Quick Align” feature in the Maestro workspace (Schrödinger Suite, Release 2022–1, Academic license) . Upon close observation of the various natural shapes of NADH, it was seen that out of 345, 259 of the 3-dimensional NADH conformations broadly fell into 6 shapes. The remaining 86 structures did not cluster into any major group. These conformations typically appeared in tiny numbers (2–3 structures per pattern) and did not form statistically or structurally meaningful clusters. These low-frequency conformations were collectively categorised as outliers (Table. S1).

As listed in the Table 1, 8 descriptors were developed: a) Internal Angle 1 (IA1), which is defined as the included angle between C4N-C1D-PN atoms, b) Internal Angle 2 (IA2), which is defined as the included angle between N1A-C1B-PA atoms, c) Internal Angle 3 (IA3), which is the included angle between C1D-PN-C1D atoms, d) Internal Angle 4 (IA4) which is the included angle between C1B-PA-C1D atoms, e) Dihedral Angle 1 (DA1) which is O4D-C1D-N1N-C2N, e) Dihedral Angle 2 (DA2) which is C4A-N9A-C1B-O4B, f) Dihedral Angle 3 (DA3) which is C4N-C3N-C7N-N7N, g) Distance 1 (D1) which is the distance between C4N-N1A and h) Distance 2 (D2) which is the distance between C1B-C1D.

**Table 1.** Naturally occurring different NADH shapes exist in various groups. (1) Group 1 (2) Group 2 (3) Group 3 (4) Group 4 (5) Group 5 (6) Group 6. Here, IA1 is the angle between C4N-C1D-PN, IA2, is the angle N1A-C1B-PA, IA3 is the angle C1D-PN-C1D, IA4 is the angle C1B-PA-C1D, DA1 is the dihedral angle between O4D-C1D-N1N-C2N, DA2 is the dihedral angle between C4A-N9A-C1B-O4B, DA3 is the dihedral angle between C4N-C3N-C7N-N7N, D1 is the distance between C4N-N1A and D2 is the distance between C1B-C1D.

| Sr No | NADH Shape | IA1 | IA2 | IA3 | IA4 | DA1 | DA2 | D1 | D2 | Number of structures with corresponding NADH shape |
| --- | --- | --- | --- | --- | --- | --- | --- | --- | --- | --- |
| 1 | Group 1 | 80-114 | 125-179 | 91-135 | 86-130 | 36-85 | -162 to -74 | 16-18 | 8-12 | 93 |
| 2 | Group 2 | 82-116 | 136-167 | 95-120 | 89-129 | -124 to -73 | -135 to -79 | 16-18 | 10-11 | 129 |
| 3 | Group 3 | 81-84 | 105-113 | 112-143 | 149-155 | -149 to -138 | -94 to -86 | 13-14 | 10-11 | 8 |
| 4 | Group 4 | 114-123 | 129-135 | 91-98 | 111-115 | 54-83 | -179 to -169 | 17-18 | 10-11 | 5 |
| 5 | Group 5 | 87-89 | 107-113 | 70-72 | 67-72 | -153.6 to -123 | -167 to -163 | 4-5 | 6-7 | 2 |
| 6 | Group 6 | 90-95 | 142-157 | 139-148 | 162-168 | -161 to -149 | -105 to -92 | 17-19 | 11-12 | 4 |

### NADH In Different Enzyme Classes

NADH was found to be co-crystallized with proteins belonging to various enzyme classes, indicating the essential and diverse functional roles of NADH. For each protein (enzyme)-ligand structure, we parsed the header record by identifying the ‘HEADER’ keyword at the beginning of each line. We then searched for the enzyme class keywords (e.g., ‘OXIDOREDUCTASE’, ‘TRANSFERASE’, ‘HYDROLASE’, ‘ISOMERASE’, ‘LIGASE’, and ‘TRANSLOCASE’, etc.) in the header record. If any of these keywords were found in the header record, we classified the protein corresponding to the respective enzyme class.

### NADH-Protein Interaction Analysis

At a high level, our computational approach to generating statistics around NAI (a three-letter identifier for NADH in the Protein Data Bank) interactions with proteins involves the extraction of the protein-NAI ligand interaction complex structures from RCSB PDB, and using Protein-Ligand Interaction Profiler (PLIP) to profile those interaction complexes corresponding to six groups of protein-NAI ligand complexes [23,25]. The PLIP reports were then programmatically parsed to generate statistics related to NAI atom interactions with the proteins in the NADH-protein complexes. A high-level schematic diagram of our computational method for generating protein-NAI ligand interaction statistics is shown in Fig.1. We used standard Python packages, Open Babel [26] and the Biopython package [27] for developing our computational method.

### PDB Files Extraction from RCSB PDB

The experimentally determined protein-ligand (NAI) complex structures (.pdb files) were extracted programmatically from RCSB PDB in batch mode [23].

### Protein-ligand Interaction Complex Profile Report Extraction

We utilized Protein-Ligand Interaction Profile (PLIP) to computationally extract protein-ligand interaction profile reports (PLIP reports) [17]. RCSB PDB files of the protein-NAI ligand complexes were input to PLIP to generate the protein-NAI ligand interaction profile reports for the corresponding protein-ligand complexes. The standard angle and distance thresholds used in our study were adopted from the PLIP, in which these parameters were shown to reliably capture the protein-ligand interactions [25]. For identifying hydrogen bonds, we used the maximum distance between the hydrogen bond donor and acceptor as 4.1Å and the minimum angle at the hydrogen bond donor as 100°. To identify hydrophobic contacts, we used a maximum distance cutoff of 4.0Å. For identifying water bridges, we used the following thresholds: minimum and maximum distances between water oxygen and polar atom as 2.5Å and 4.1Å respectively, minimum and maximum angles between acceptor, water oxygen, and donor hydrogen as 71° and 140° respectively, and minimum angle between water oxygen, donor hydrogen and donor atom as 100°.

### B-factor Extraction Corresponding to the Protein-Ligand Interaction Complex

We parsed the protein-ligand interaction profile reports (PLIP report) to identify the protein residue coordinates corresponding to protein-NAI ligand interactions. We then parsed the corresponding PDB coordinate files to identify the B-factors corresponding to each protein residue participating in hydrophobic interactions, hydrogen bonds, and water bridges, and generated statistics (average range and distribution) at the aggregate level for the B-factors of each group of protein-NAI ligand complexes (Fig. 1).

**Figure 1:**
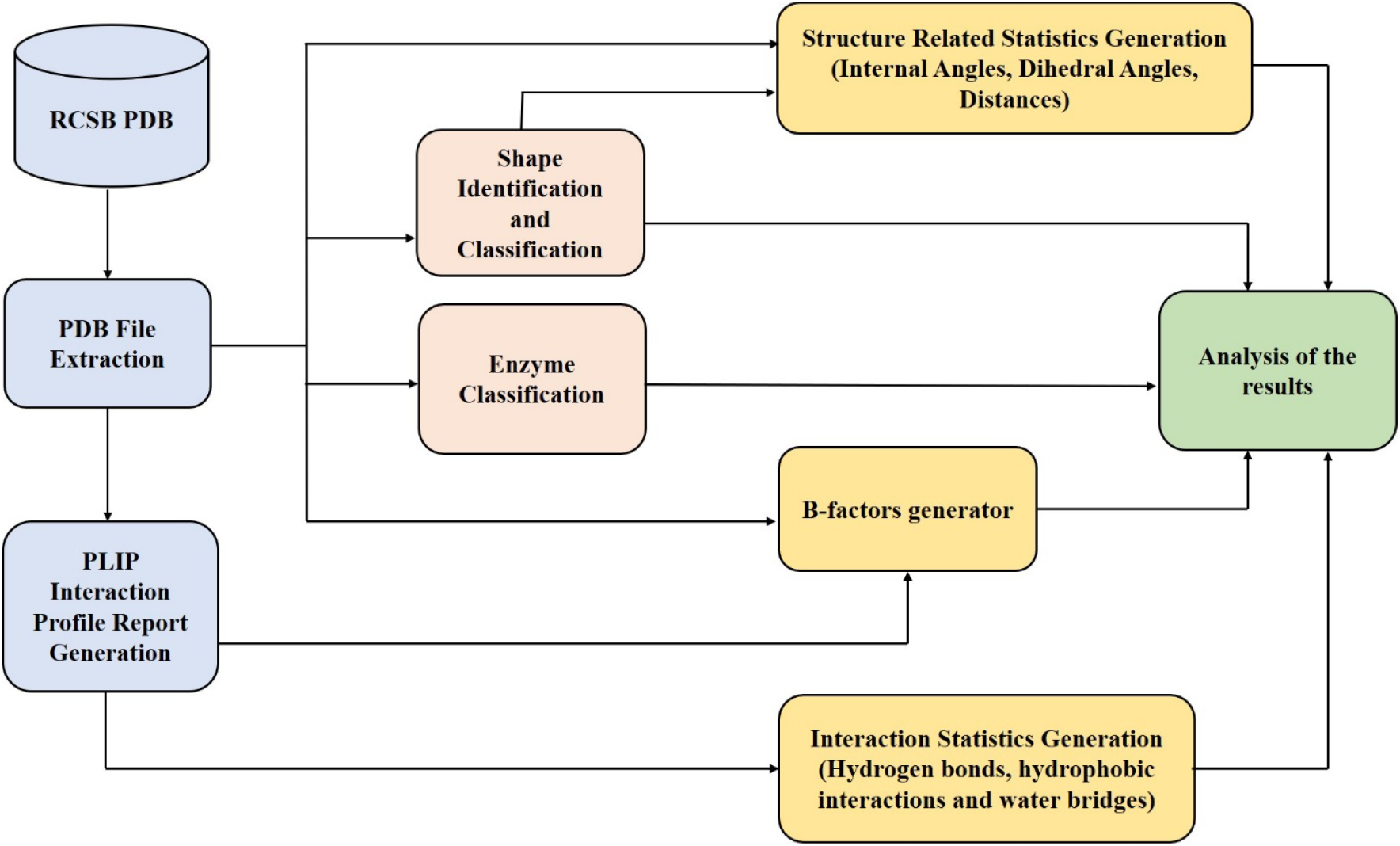
A schematic diagram of our programmatic approach for generating protein-NAI ligand complex leveraging RCSB PDB and PLIP.

### Generation of Statistics

We then computationally parsed each protein-NAI ligand interaction complex file (extracted from RCSB PDB) to generate following structure related statistics around NAI atoms: (1) Internal angles between (a) (C4N, C1D, PN) atoms called IA1, (b) (N1A, C1B, PA) atoms called IA2, (c) (C1D, PN, C1B) atoms called IA3, and (C1B-PA-C1D) called IA4 (2) dihedral angle between (O4D, C1D, N1N, C2N) atoms called DA1 and dihedral angle between (C4A, N9A, C1B, O4B) atoms called DA2, and (3) distances between (a) (C4N, N1A) atoms (D1), and (b) (C1B, C1D) atoms (D2). We also computationally identified the NAI atoms that are participating in hydrogen bonds, hydrophobic interactions and water bridges against each protein-NAI ligand complex corresponding to each six-group of interaction complexes and their distribution across the 20 standard amino acids.

## Results

### Natural NADH Shapes And Structural Plasticity

The superposition of all the NADH coordinates broadly showed NADH falling into six distinct shapes, which were named Group 1 - Group 6 (Table 1 and Fig. 2). The shape of NADH in groups 1 and 2 seems to be nature’s preferred shape, as they represent 65.2% of the total NADH PDB dataset. These two groups share similar D1 and D2 values, consistent with comparable compaction of the Adenine and Nicotinamide rings, but differ systematically in DA1 and DA2 sign and range, implying alternative torsional states of the pyrophosphate. Groups 3 and 4 are rare (8 and 5 structures, respectively) and exhibit narrower ranges for IA1–IA3 but distinct dihedral signatures. Groups 5 and 6 are extremely under-represented (2 and 4 structures) and exhibit both smaller D1 values and more extreme DA1/DA2 ranges. D1 (C4N–N1A) remains within a restricted range in Groups 1–4, consistent with a relatively conserved separation between the adenine and nicotinamide rings that supports efficient electron transfer and stacking interactions with catalytic residues. In contrast, the smaller D1 in Group 5 and the slightly larger D1 range in Group 6 indicate compacted or expanded cofactor shapes, respectively, which could modulate hydride transfer distance or alter how the nicotinamide ring aligns with substrate.

**Figure 2.**
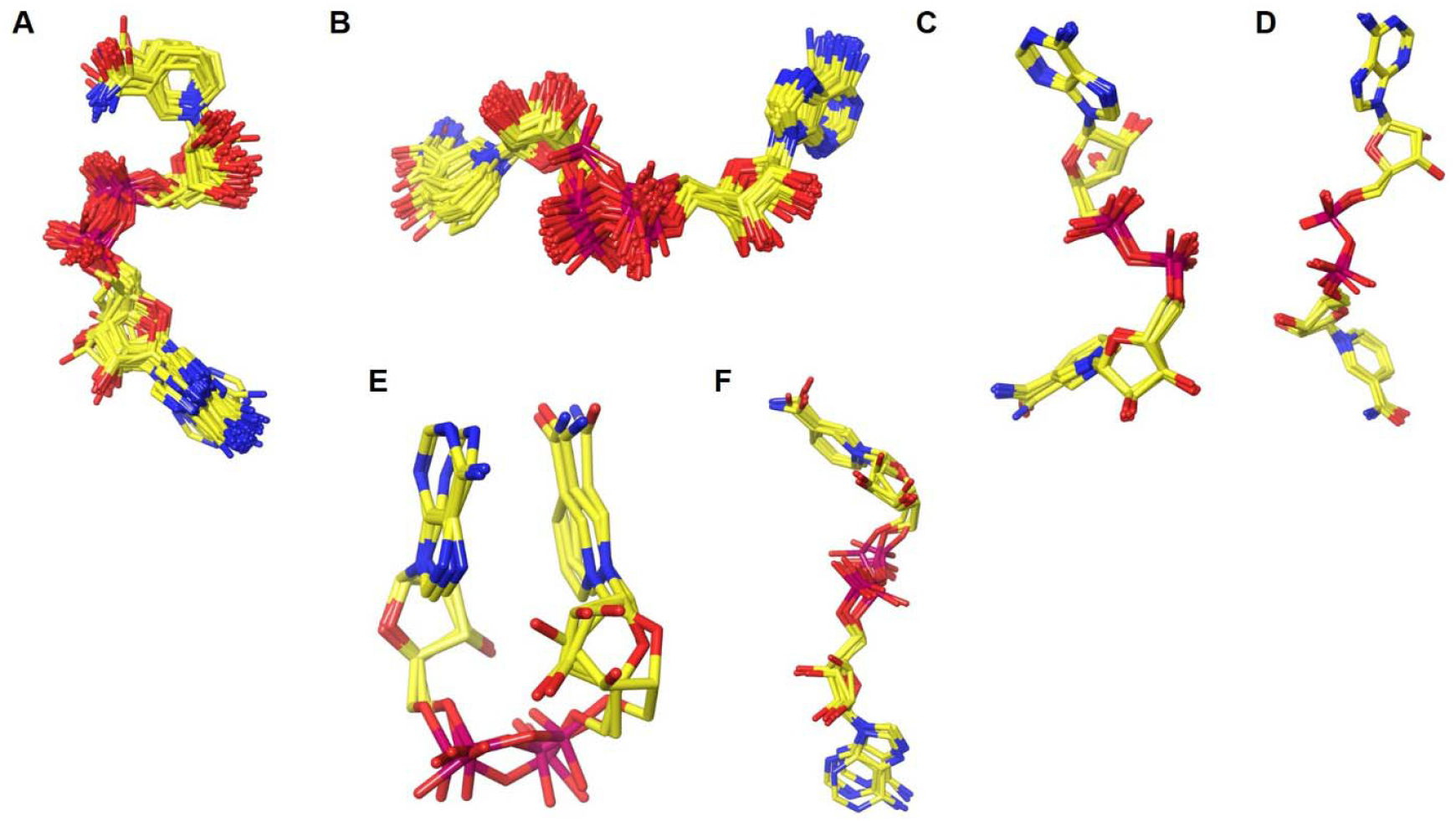
Naturally occurring shapes of NADH. (A) Group 1 (B) Group 2 (C) Group 3 (D) Group 4 (E) Group 5 (F) Group 6. Refer to Table 1 for the description of these groups.

When the space in the NADH binding site in the respective proteins was correlated with the shapes of the NADH, it was clear that NADH could achieve its hydride transfer task by adapting and adjusting to the available space and surrounding environment (Fig. 3).

**Figure 3.**
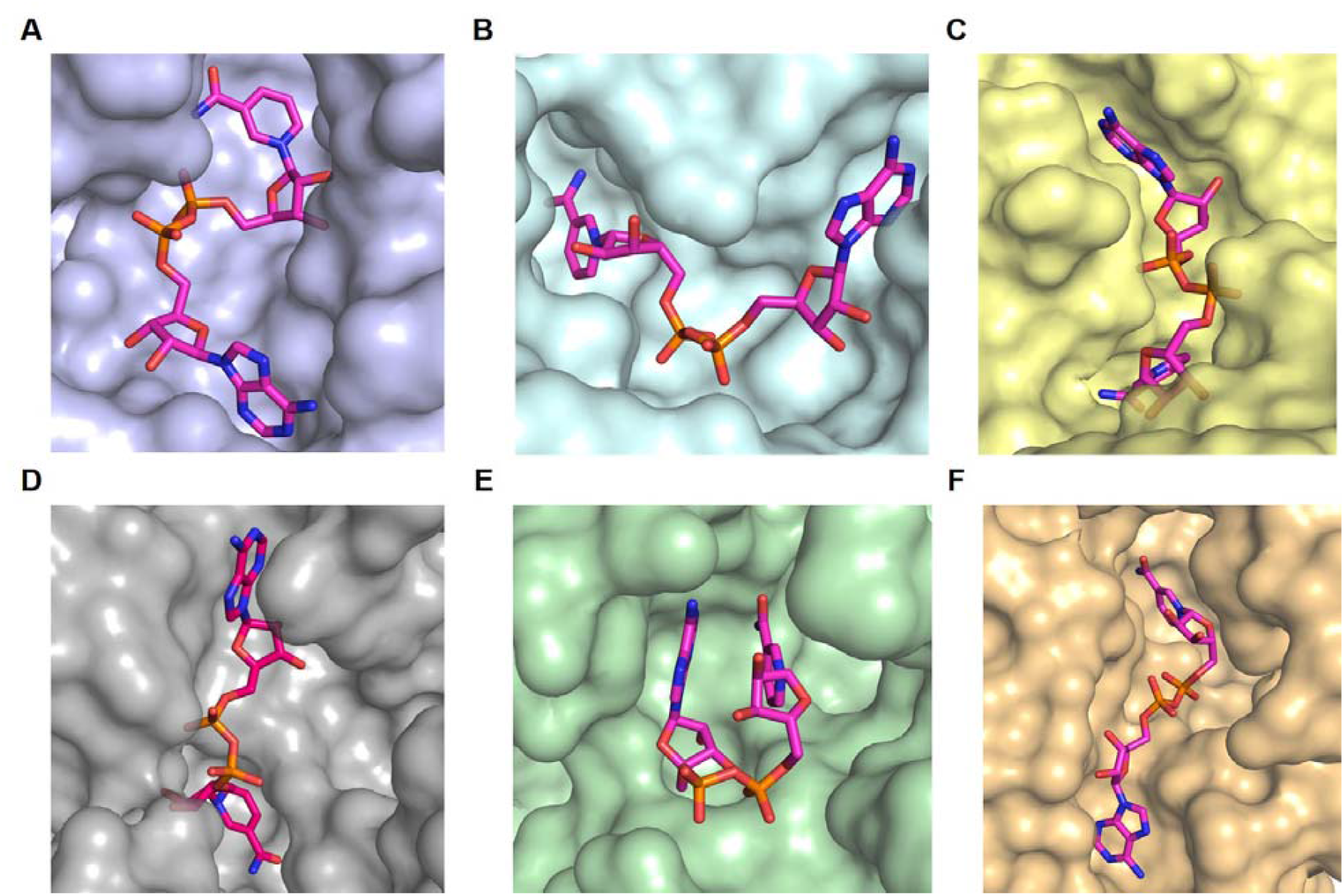
The NADH binding pocket and shape complementary. (A) Group 1 (B) Group 2 Group 3 (D) Group 4 (E) Group 5 (F) Group 6

### NADH Interactions In Different Classes Of Enzymes

NADH-bound structures are overwhelmingly associated with oxidoreductases across all the groups, with only a small fraction mapping to other classes. Fig 4A shows oxidoreductases account for almost all NADH structures in the dataset, whereas the other classes, which are translocases, hydrolases, isomerases, transferases, and lyases, contribute to only a handful of entries each. Group 1 and 2 (Fig. 4B and 4C) comprise the major NADH conformational cluster, which is also dominated by oxidoreductases.

**Figure 4.**
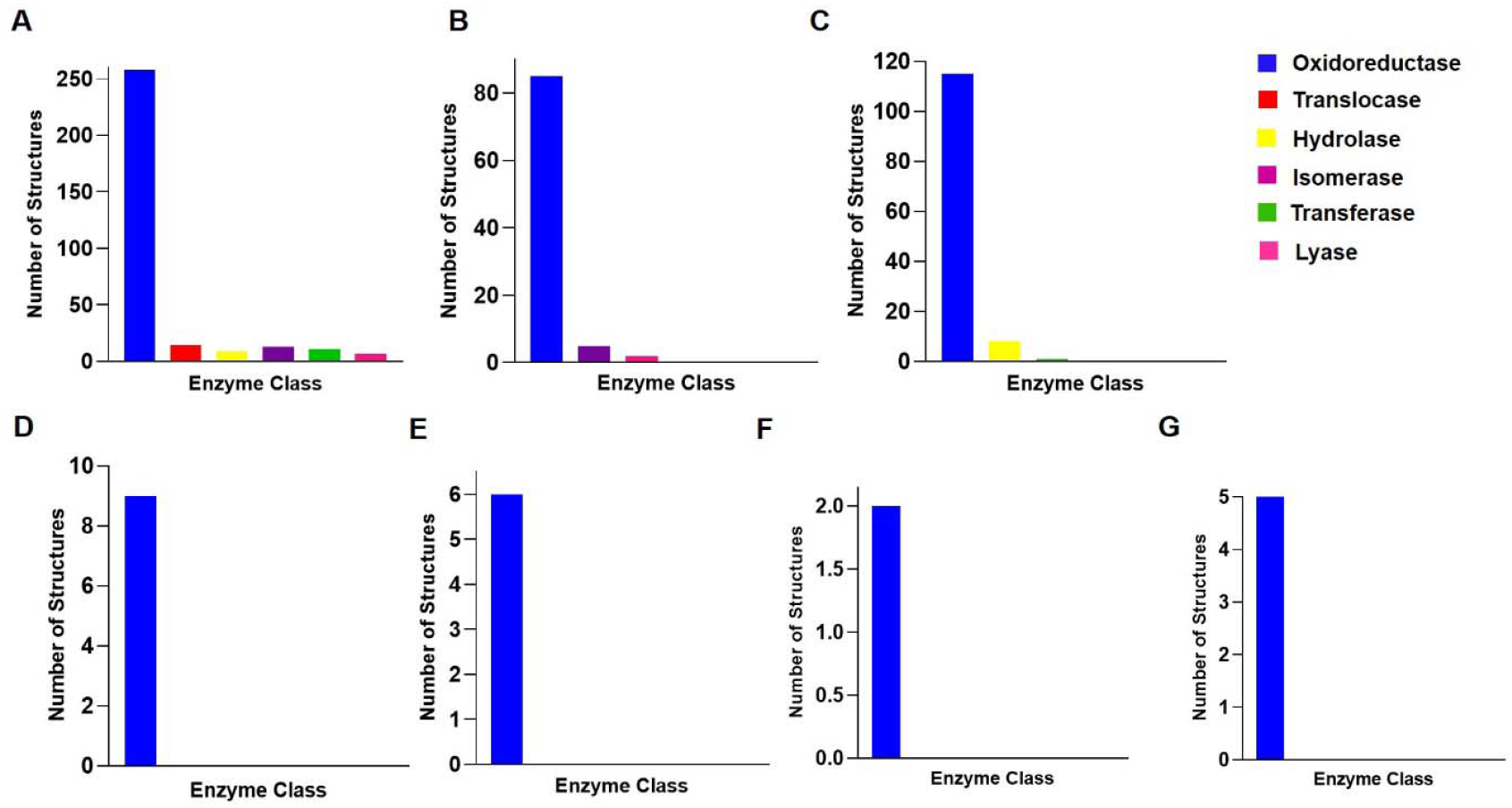
Enzyme classes of NADH amongst all groups. (A) Overall (B) Group 1 (C) Group 2 (D) Group 3 (E) Group 4 (F) Group 5 (G) Group 6

### NADH-Protein Atomic Interactions And B-factor Analysis

The results of NADH interacting and non-interacting atoms are shown in Fig. 5B. The table shows that almost all heteroatoms of NADH participate in protein interactions. In contrast, only a small subset of carbons are directly involved in the interactions. All seven nitrogen atoms and all fourteen oxygen atoms are classified as interacting, indicating that proteins for hydrogen bonding, electrostatic interactions, or metal coordination exploit every N and O position on NADH. The two phosphate phosphorus atoms (PA, PN) are listed as non-interacting despite being charged. Here, the side chain and backbone amide hydrogens bind to the oxygens of the phosphate, whereas the P center remains sterically shielded. Out of 21 carbons, only three (C3N, C4N, C5N) are marked as interacting. Interestingly, all of them are located in the nicotinamide ribose region. The remaining 18 carbons, distributed over the adenine ribose, nicotinamide ring periphery, and linker region, are non-interacting. The NADH binding is predominantly driven by its N and O atoms through polar contacts.

**Figure 5.**
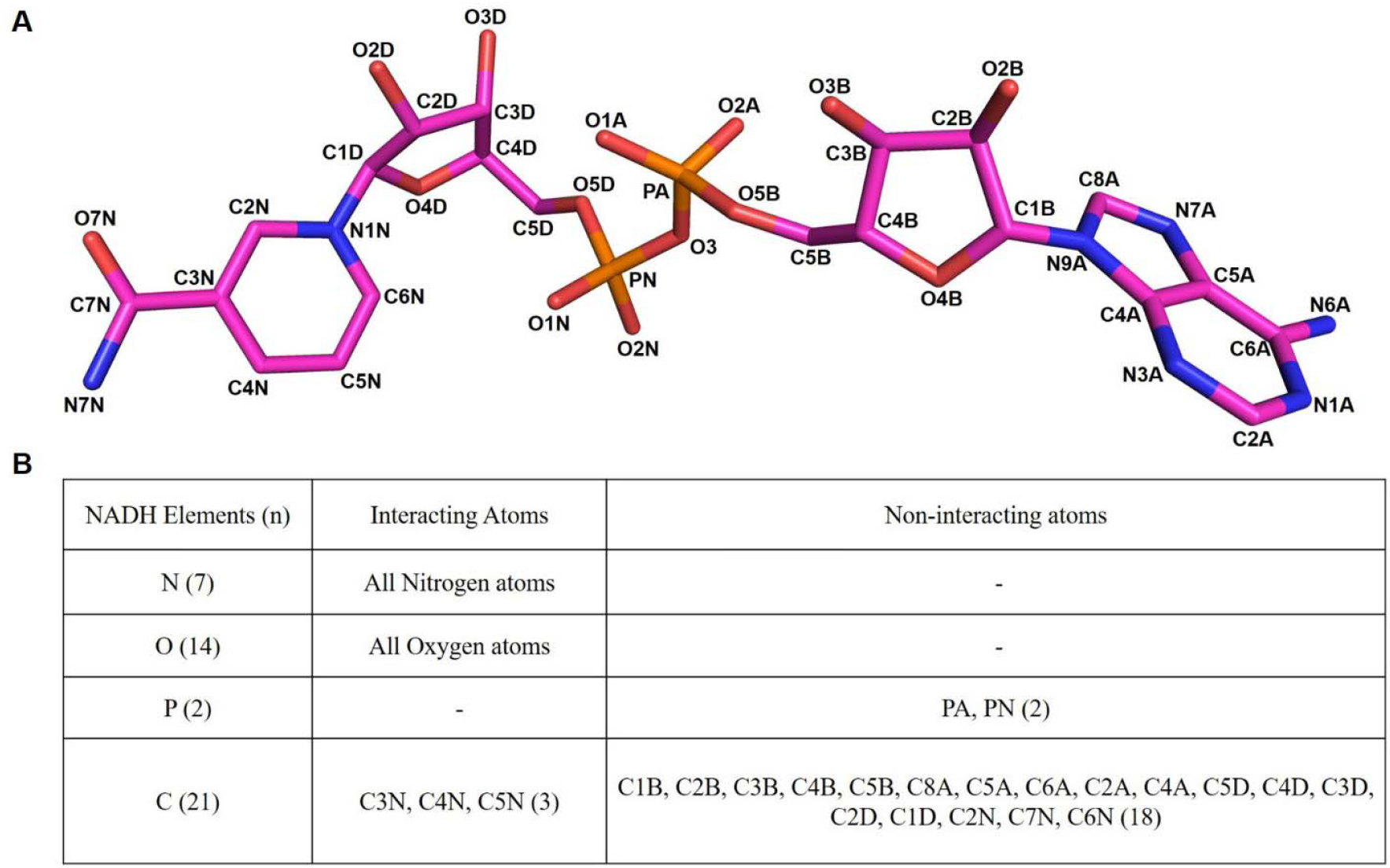
NADH-protein atomic interactions. (A) NADH naming and numbering conventions. Standard atom names are per PDB format. (B) NADH interacting and non-interacting atoms (n = number of atoms).

The B-factor distributions across the six NADH groups are shown in (Fig. S1). It highlights the relation between the distinct binding modes and the ligand stability. While groups 1 and 2 have the largest number of NADH-bound structures, they also have the widest B-factor ranges. These indicate substantial variability in NADH mobility across proteins within these groups. Despite their large range, the high number of these groups suggests that flexible NADH binding modes are widely tolerated and evolutionarily preferred across enzyme classes. In contrast, groups 4 and 5 display narrow B-factor ranges and a rigid NADH binding; however, their limited number indicates that such constrained binding modes are less common and are likely limited to specific enzymes. Groups 3 and 6 reflect intermediate B-factor spreads.

### NADH-Protein Residue Interaction Analysis

To identify conserved residue preferences around NADH, we mapped the interaction frequency of each amino acid with every atom of NADH across all structures in our dataset. The residue-level interaction map summarises the most common residues contacting each atom of NADH (Fig. 6). The adenine moiety is most frequently contacted by residues such as Ile at its nitrogen atoms, while Arg and Asp interact with the exocyclic amine (NH2). The adjacent ribose shows prominent contacts with Glu, Asp, and Leu, especially at hydroxyl groups, indicating a preference for polar and hydrophobic residues in stabilising these positions. Ala appears frequently at the phosphate-ribose linkage. Phosphate oxygens within the pyrophosphate region commonly interact with Gly, Arg, and Ser, demonstrating a recurring involvement of small and basic residues at sites of negative charge. The nicotinamide ring interacts most frequently with Glu, Thr, Gly, Asn, Ser, Val, and His, reflecting the chemical variability of the aromatic ring and its amide and carboxamide functionalities. The terminal amide group interacts strongly with Glu, Thr, Gly, and Asn at both its nitrogen and oxygen atoms, indicating a motif for hydrogen bonding and charge interaction stabilisation. The ribose attached to nicotinamide shows common interactions with Asn, Ile, Thr, and Val at its hydroxyl sites.

**Figure 6.**
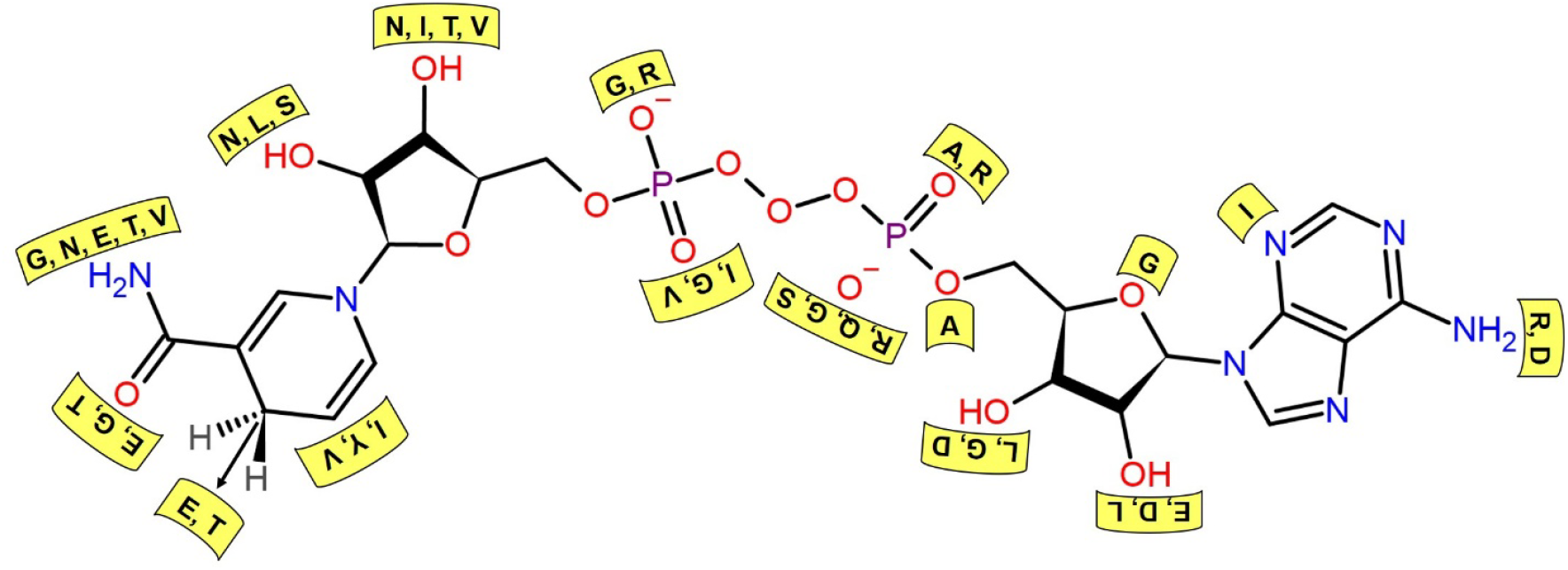
Residue-level interactions. The figure shows the most frequently observed protein residues contacting each atom of NADH across all analyzed structures. The protein residues are shown in yellow boxes while the atoms of NADH, which are O, N, H and P, are shown in red, blue, grey and purple colors, respectively.

### NADH Moiety Level Interactions

To understand how different regions of the NADH contribute to the binding and ligand recognition across diverse protein structures, we studied moiety-specific interaction frequencies in NADH. We divided the NADH structure into three moieties. The Nicotinamide moiety consists of the nicotinamide ring, the ribose sugar attached to it, the next is the adenine moiety, which has the adenine ring and the corresponding ribose sugar, and the last moiety is the phosphate moiety which is a pyrophosphate group in the middle of both the nicotinamide and the adenine moieties. The atomic distribution of all three moieties is shown in Table 2.

**Table 2.** Atomic-level distribution. Atomic distribution of all three moieties of NADH

| Sr No | Moiety | Corresponding Atoms |
| --- | --- | --- |
| 1 | Nicotinamide | C5D, C4D, O4D, C3D, O3D, C2D, O2D, C1D, N1N, C2N, C3N, C7N, O7N, N7N, C4N, C5N, C6N |
| 2 | Phosphate | O3, PN, O1N, O2N, O5D, PA, O1A, O2A, O5B |
| 3 | Adenine | C5B, C4B, O4B, C3B, O3B, C2B, O2B, C1B, N9A, C8A, N7A, C5A, C6A, N6A, N1A, C2A, N3A, C4A |

Across all the groups, the nicotinamide moiety seems to be the predominant interaction hotspot. In group 1 (Fig 7B), the interaction profile displays an almost balanced distribution among all three moieties with the nicotinamide moiety contributing the largest share (38.46%), followed by the phosphate moiety (32.01%) and then the adenine moiety (29.5%). A similar pattern is observed in group 2 (Fig 7C) where among the three moieties, the nicotinamide moiety interactions are dominant compared to the other two moieties. In group 3 (Fig. 7D), there is a marked shift towards nicotinamide-centric interactions. The nicotinamide moiety interactions in group 3 are 59.87%, followed by adenine moiety with 22.22% and then phosphate moiety with 17.90%. In contrast to the earlier groups, adenine and nicotinamide moiety interactions show almost equal contribution in group 4 (Fig 7E) with 42.85% of adenine and 40.47% of nicotinamide interactions. In group 5 (Fig 7F), the adenine accounts for the largest share with 50% of the total interactions. Whereas, nicotinamide and phosphate constituted 29.54% and 20.45% of total interactions, respectively. Group 6 (Fig 7G) returns to the balanced interaction profile with a moderate contribution from all three moieties.

**Figure 7.**
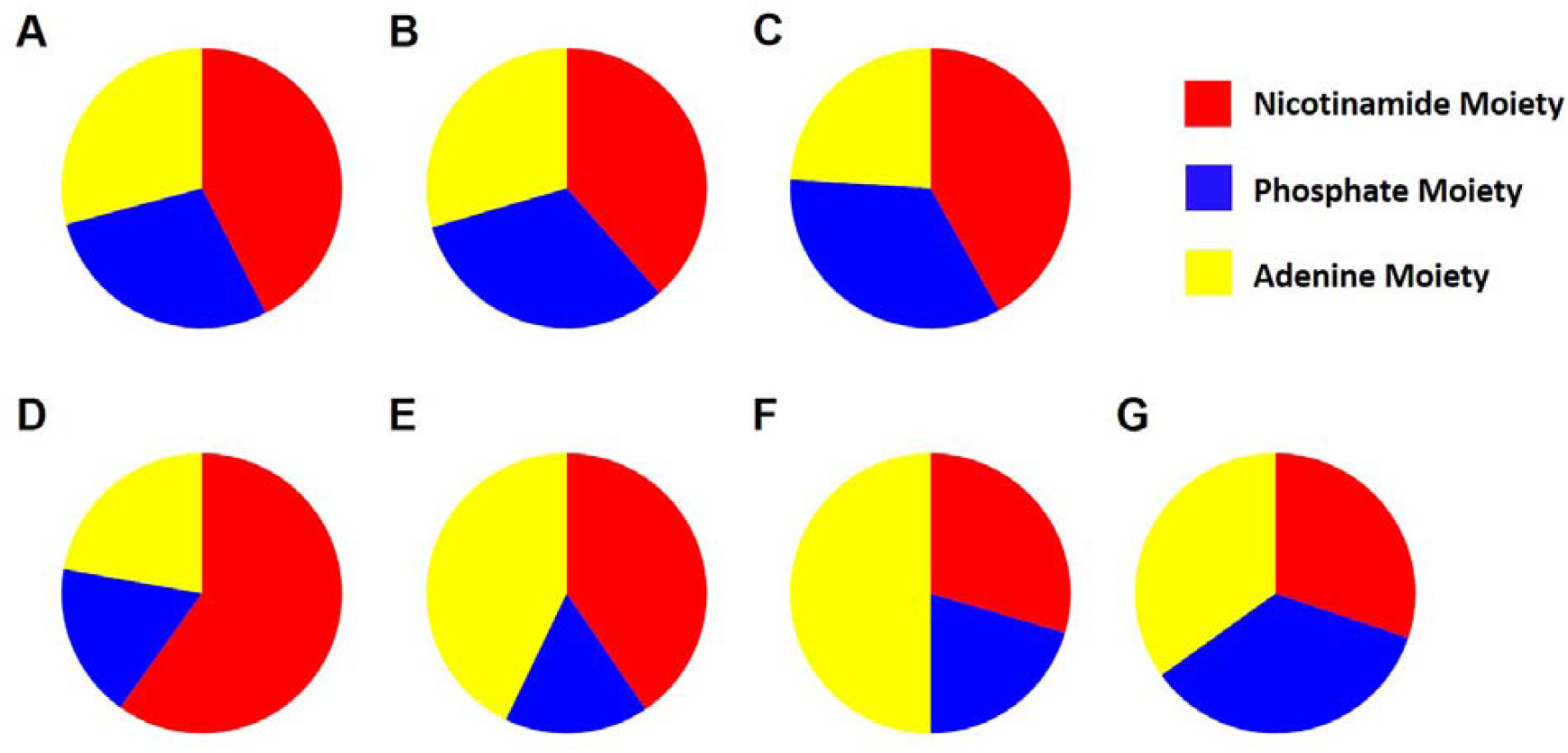
Moiety level interactions. Moiety level interactions of of NADH amongst all groups (A) Overall (B) Group 1 (C) Group 2 (D) Group 3 (E) Group 4 (F) Group 5 (G) Group 6

## Discussion

The systematic analysis of 345 NADH-bound crystal structures reveals that the NADH populates a set of six recurrent 3D shapes (Fig. 2). The strong enrichment of Groups 1 and 2, which together represent almost two-thirds of the dataset, suggests that nature preferentially chooses conformers in which the adenine and nicotinamide rings maintain a conserved separation. At the same time, the pyrophosphate adapts to only a few torsional states. This supports the view that efficient hydride transfer and interaction with the catalytic residues require both spatial proximity of the redox centers and constrained orientation of the linker.

The rarity of Groups 3–6 indicates that more compacted or expanded NADH shapes are tolerated in only a handful of binding pockets and may correspond to specialised catalytic strategies or regulatory states. In particular, the smaller D1 values in Group 5 and the larger D1 range in Group 6 imply altered hydride transfer distances or changes in how the nicotinamide ring aligns with adenine, substrates, and active-site residues (Table. 1). These unusual conformations offer interesting targets for engineering enzymes or designing selective inhibitors that exploit typical cofactor geometries.

The enzyme class distribution highlights the centrality of NADH to oxidoreductases, which account for most of the complexes, while other EC classes contribute only a small proportion of structures (Fig. 4). This reflects the canonical redox role of NADH but also highlights underexplored interfaces in non-oxidoreductase families.

At the atomic level, the interaction analysis shows that nearly all heteroatoms of NADH are utilised by proteins. In contrast, only a small subset of carbons—concentrated in the nicotinamide ribose region—participates directly in contacts (Fig. 5). This pattern suggests that NADH recognition is primarily driven by hydrogen bonding and electrostatic interactions with N and O atoms. At the same time, the majority of the carbon skeleton serves as a structurally supportive but chemically inert scaffold; meanwhile, the two pyrophosphate atoms remain non-interacting. The B-factor analysis suggests that while rigid NADH binding confers high local stability, flexible binding modes dominate across the NADH-dependent enzyme landscape (Fig. S1).

The residue-level interaction map further clarifies how NADH interacts with the protein surface (Fig. 6). Adenine atoms are often contacted by charged and hydrophobic residues such as Arg, Asp and Ile. A similar trend is observed in the adjacent ribose hydroxyls, where charged and non-polar residues dominate the interactions. The pyrophosphate oxygens frequently engage with Gly, Ser, and Arg, consistent with a requirement for small backbone donors and basic side chains to neutralise negative charge. The nicotinamide moiety interacts with a chemically diverse set of residues, including Glu, Thr, Gly, Asn, Ser, Val, and His, highlighting its dual role as a redox centre and a versatile hydrogen-bond donor/acceptor platform.

Moiety-level analysis shows that the nicotinamide region is the primary interaction hotspot in most groups, particularly in Group 3, where it accounts for nearly 60% of contacts (Fig. 7). In contrast, Group 5 displays adenine-centered binding, suggesting that some rare conformers rely on adenine-driven recognition while keeping nicotinamide more solvent-exposed or mobile. Such differences imply that the same cofactor can be recognised through multiple interaction modes.

The identification of non-interacting carbon positions across the NADH scaffold suggests chemically tolerant sites where modifications are less likely to disrupt the binding pose or catalytic geometry. In contrast, substitutions at interacting heteroatoms or at the nicotinamide carbons that make frequent contacts would be expected to impair affinity or alter the orientation required for hydride transfer. Together, the shape clustering, atom-level contact maps, and moiety-level preferences provide a structural framework for rationally designing NADH analogues with tuned binding specificity, altered redox properties, or selective inhibition of a particular enzyme subclass.

Conformational flexibility of ligands or lead compounds, due to rotatable single bonds, hinders their development as nanomolar binding affinity drugs/inhibitors against drug targets because of the flexibility that comes with an entropy cost in the free energy of binding (ΔG_bind_). These types of analyses of ligand or cofactor conformations in the target molecules would help to fix the shape of a ligand to a single or a few conformations through the introduction of double or triple bonds, or ring strictures, so that reduction of conformational freedoms dramatically enhances the binding affinity by reducing the entropy cost to ΔG_bind_.

## Conclusion

This work provides a global, structure-based view of how NADH is recognised across diverse protein environments. By mining 345 co-crystal structures, six preferred NADH shapes were identified, with two dominant conformers that preserve a conserved adenine–nicotinamide separation while sampling only a limited set of pyrophosphate torsions. Atom- and residue-level analyses showed that virtually all heteroatoms of NADH are exploited for binding, whereas only a few carbons—mainly in the nicotinamide ribose region—serve as direct contact points.

Moiety-level interaction profiling revealed that the nicotinamide segment is the principal hotspot in most complexes, although some rare conformational groups display adenine-centric or more balanced interaction patterns. The structural descriptors and interaction maps reported here offer practical guidelines for cofactor engineering and for designing NADH-mimetic inhibitors that selectively target specific NADH-dependent enzymes.

## Supporting information

https://docs.google.com/document/d/1gt4t8sVpi3TezrJm2dtisyz9ZjIPfOei23c1Lz5om8k/edit?usp=sharing

## Acknowledgements

The authors would like to thank IITH for intramural funding. D.S. is supported by the Council of Scientific and Industrial Research, India.

## Appendix A

Supplementary Information

## Research Data

Code and Data

