## Supplementary material for "Conformational Diversity and Interaction Signatures of NADH across protein families": https://docs.google.com/document/d/1gt4t8sVpi3TezrJm2dtisyz9ZjIPfOei23c1Lz5om8k/edit?usp=sharing

| 1GIQ | 2DLD | 2YVJ | 3QVT | 4ILK | 4YAC | 5ZJ5 | 6SBU | 7Q3A | 8YQ3 |
| --- | --- | --- | --- | --- | --- | --- | --- | --- | --- |
| 1OG4 | 2FRD | 3AFO | 3RSQ | 4JAF | 4YB4 | 6I1P | 6TF0 | 7WMZ | 9CP7 |
| 1LSU | 2GR9 | 3FHM | 3RTD | 4LTN | 4YKF | 6KE3 | 6VW7 | 8AD4 | 9CP8 |
| 1N2S | 2GWL | 3FLK | 3TE5 | 4P53 | 4ZCC | 6MVT | 7BR4 | 8AD5 | 9E4U |
| 1VRW | 2HMS | 3FMX | 4FJU | 4PLV | 5VWU | 6Q9C | 7CTL | 8CT0 | 9FDJ |
| 1NXG | 2HMT | 3IAM | 4FW8 | 4R69 | 5WU0 | 6Q9G | 7JSO | 8SWY | 9FE5 |
| 1WNB | 2IMP | 3IQD | 4G6H | 4XSH | 5YIJ | 6Q9J | 7MJ1 | 8V36 | 9FIF |
| 1O0S | 2J6L | 3PRJ | 4H8A | 4Y9D | 5YVT | 6Q9K | 7MYC | 8WFQ | 9FIH |
| 9H4X | 9IVY | 5BWM | 1L7E | 5H04 | 5TLB |  |  |  |  |

**Table S1. List of outliers**. These structures could not be classified into any shape.

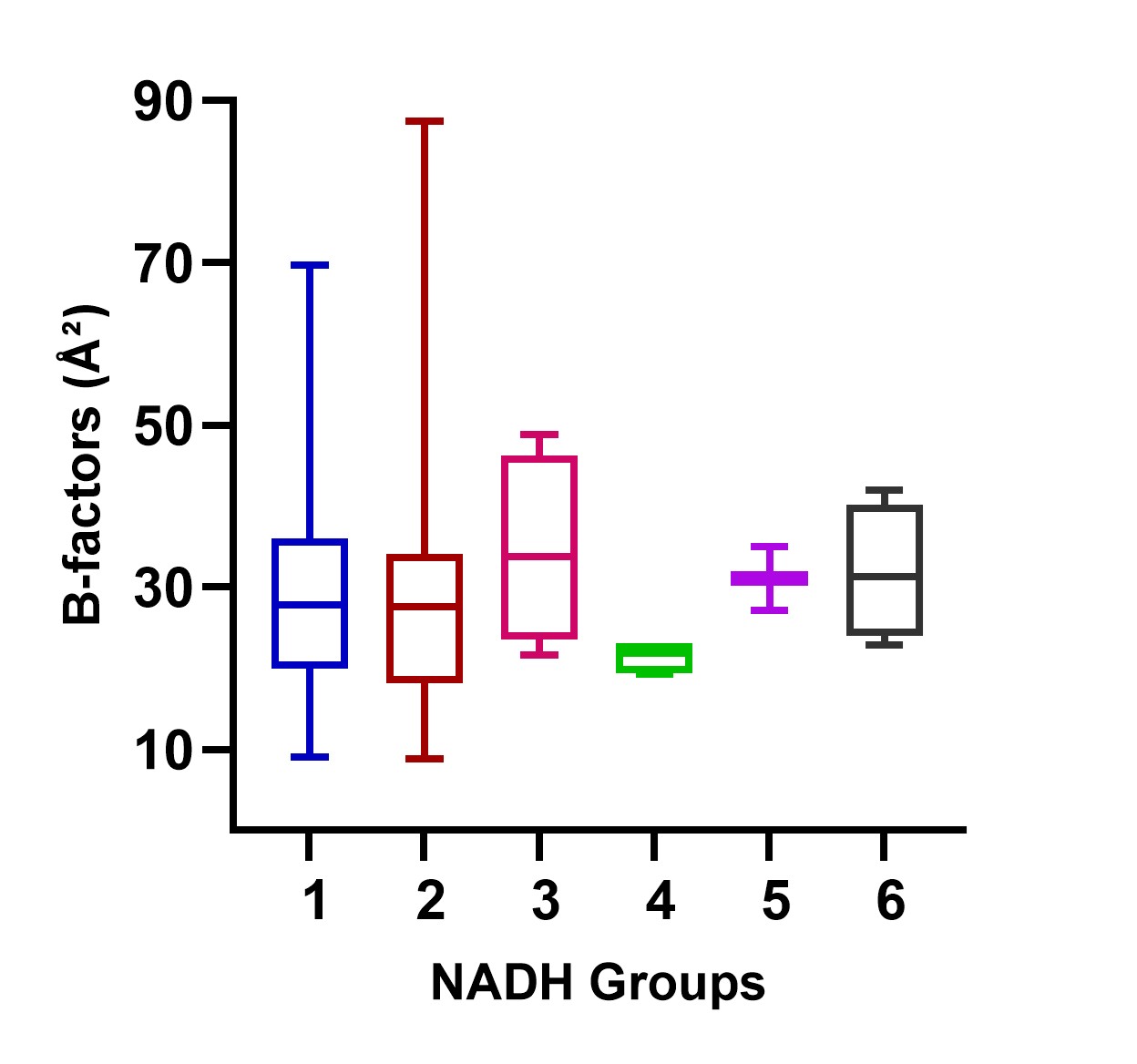

**Figure S1**. **The distribution of NADH B-factors across six groups.** The central line represents the median, boxes denote the interquartile range, and whiskers indicate the spread of the data. Groups 1 and 2 display broad B-factor distributions, while Groups 4 and 5 exhibit narrow B-factor distributions. Groups 3 and 6 display intermediate distributions, characterised by moderate interquartile ranges.
